# Decoding face recognition abilities in the human brain

**DOI:** 10.1101/2022.03.19.484245

**Authors:** Simon Faghel-Soubeyrand, Meike Ramon, Eva Bamps, Matteo Zoia, Jessica Woodhams, Anne-Raphaelle Richoz, Roberto Caldara, Frédéric Gosselin, Ian Charest

**Author notes:** Corresponding authors: Ian Charest, Simon Faghel-Soubeyrand.

## Abstract

Why are some individuals better at recognising faces? Uncovering the neural mechanisms supporting face recognition ability has proven elusive. To tackle this challenge, we used a multi-modal data-driven approach combining neuroimaging, computational modelling, and behavioural tests. We recorded the high-density electroencephalographic brain activity of individuals with extraordinary face recognition abilities—super-recognisers—and typical recognisers in response to diverse visual stimuli. Using multivariate pattern analyses, we decoded face recognition abilities from 1 second of brain activity with up to 80% accuracy. To better understand the mechanisms subtending this decoding, we compared computations in the brains of our participants with those in artificial neural network models of vision and semantics, as well as with those involved in human judgments of shape and meaning similarity. Compared to typical recognisers, we found stronger associations between early brain computations of super-recognisers and mid-level computations of vision models as well as shape similarity judgments. Moreover, we found stronger associations between late brain representations of super-recognisers and computations of the artificial semantic model as well as meaning similarity judgments. Overall, these results indicate that important individual variations in brain processing, including neural computations extending beyond purely visual processes, support differences in face recognition abilities. They provide the first empirical evidence for an association between semantic computations and face recognition abilities. We believe that such multi-modal data-driven approaches will likely play a critical role in further revealing the complex nature of idiosyncratic face recognition in the human brain.

**Significance:** The ability to robustly recognise faces is crucial to our success as social beings. Yet, we still know little about the brain mechanisms allowing some individuals to excel at face recognition. This study builds on a sizeable neural dataset measuring the brain activity of individuals with extraordinary face recognition abilities—super-recognisers—to tackle this challenge. Using state-of-the-art computational methods, we show robust prediction of face recognition abilities in single individuals from a mere second of brain activity, and revealed specific brain computations supporting individual differences in face recognition ability. Doing so, we provide direct empirical evidence for an association between semantic computations and face recognition abilities in the human brain—a key component of prominent face recognition models.

## Introduction

The ability to robustly recognise the faces of our colleagues, friends and family members is paramount to our success as social beings. Our brains complete this feat with apparent ease and speed, in a series of computations unfolding within tens of milliseconds in a wide brain network comprising the inferior occipital gyrus, the fusiform gyrus, the superior temporal sulcus, and more anterior areas such as the anterior temporal lobe (Duchaine & Yovel, 2015; Grill-Spector et al., 2017; Kanwisher et al., 1997). Accumulating neuropsychological and behavioural evidence indicates that not all individuals, however, are equally competent at recognising faces in their surroundings (White and Burton 2022). Developmental prosopagnosics show a great difficulty at this task despite an absence of brain injury (Susilo & Duchaine, 2013). In contrast, super-recognisers exhibit remarkable abilities for processing facial identity, and can recognize individuals even after little exposure several years before (Noyes et al., 2017; Ramon, 2021; Russell et al., 2009). The specific nature of the neural processes responsible for these individual differences remains largely unknown.

So far, individual differences studies have used univariate techniques to investigate face-specific aspects of brain processing. This revealed that contrasts between responses to faces compared to non-faces, measured by the N170 event-related potential component or by the blood oxygen level dependent signals in regions of interest, are modulated by ability (Elbich and Scherf 2017; Herzmann et al. 2010; Huang et al. 2014; Kaltwasser et al. 2014; Lohse et al. 2016; Rossion et al. 2020; Nowparast Rostami et al. 2017). However, univariate and contrast approaches are limited in their capacity to reveal the precise nature of the underlying brain computations (Vinken et al. 2022; Visconti di Oleggio Castello et al. 2021; Dwivedi et al. 2021; Harel et al. 2013).

Here, we tackled this challenge with a data-driven approach. We examined the functional differences between the brains of super-recognisers and typical recognisers using decoding and representational similarity analyses (RSA; Kriegeskorte et al. 2008; Kriegeskorte and Kievit 2013; Charest et al. 2014; Dwivedi et al. 2021; Kriegeskorte and Diedrichsen 2019) applied to high-density electrophysiological (EEG) signals and artificial neural network models. We recruited 33 participants, including 16 super-recognisers, i.e., individuals better than the 98th percentile on a battery of face recognition tests (Russell et al., 2009; **Fig. 1a**). We measured EEG in more than 100,000 trials while participants performed a one-back task. The objects depicted in the stimuli belonged to multiple visual categories including face images of different sexes, emotions, and identities, as well as images of man-made and non-face natural objects (e.g., a computer, a plant), animals (e.g., a giraffe, a monkey), and scenes (e.g., a city, a dining room; **Fig. 1b**).

**Figure 1.**
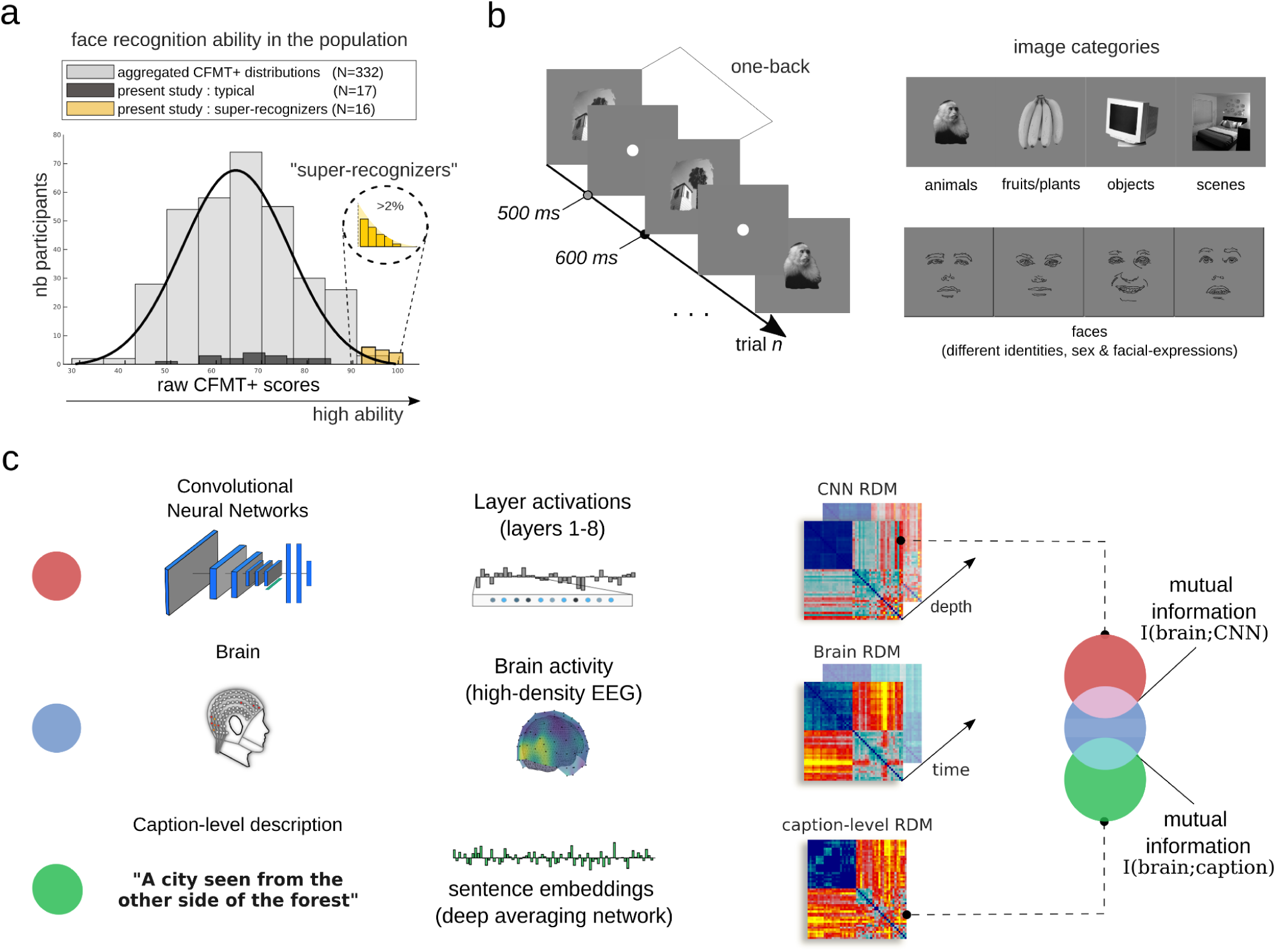
Experimental procedure and computational modelling of brain representations. **a)** The histogram shows the Cambridge Face Memory Test long-form (CFMT+, Russell et al. 2009) scores of super-recognisers (yellow bars), typical recognisers (black bars), and an additional 332 neurotypical observers from three independent studies for comparison (Faghel-Soubeyrand et al., 2019; Fysh et al., 2020; Tardif et al., 2019). **b)** Participants engaged in a one-back task while their brain activity was recorded with high-density electroencephalography. The objects depicted in the stimuli belonged to various categories, such as faces, objects, and scenes. Note that the face drawings shown here are an anonymised substitute to the experimental face stimuli presented to our participants. **c)** Representational dissimilarity matrices (RDM) were computed from convolutional neural networks (CNN; Krizhevsky et al., 2012; Simonyan & Zisserman, 2014) of vision, human brain activity, and a deep neural network of caption classification and sentence semantics (Cer et al., 2018). To characterise the CNN RDMs, we computed the pairwise similarity between unit activation patterns for each image independently in each CNN layer. The caption-level RDMs were derived from human caption descriptions of the images transformed into sentence embeddings. Brain RDMs were computed using cross-validated decoding performance between the EEG topographies from each pair of stimuli at every 4 ms time-point. Mutual information (Ince et al., 2017) between the model RDMs and the brain RDMs was assessed, for every participant, at each 4 ms step from stimulus onset.

## Results

### Discriminating super-recognisers and typical recognisers from 1 second of brain activity

With this sizable and category-rich dataset, we first attempted to classify a participant as either a super- or a typical recogniser based solely on their brain activity. More specifically, we trained Fisher linear discriminants to predict group membership from single, 1-second trials of EEG patterns (in a moving searchlight of five neighbouring electrodes). We observed up to ∼80% cross-validated decoding performance, peaking over electrodes in the right hemisphere. This performance is impressive given that the noise ceiling imposed on our classification by the test-retest reliability of the CFMT+ (Duchaine and Nakayama 2006; Russell et al. 2009), the gold-standard test used to identify super-recogniser individuals, is ∼93% (SD=2.28%; see Supplementary material). To reveal the time course of these functional differences, we applied the same decoding procedure to each 4-ms interval of EEG recordings. Group-membership predictions attained statistical significance (*p*<.001, permutation tests, **Fig. 2a**) from about 65 ms to at least 1 s after stimulus onset, peaking around 135 ms, within the N170 window (Bentin et al., 1996; Rossion & Jacques, 2012). Notably, similar results were obtained following the presentation of both face *and* non-face visual stimuli (**Fig. 2a; see also Supplementary Fig. 1**). The decoding of group-membership from non-face stimuli could be due to face features stored in short-term memory from one-back trials. To control for this possibility, we repeated our decoding analysis for non-face trials either preceded by face trials or by non-face trials. We found significant decoding of group membership in both cases (**Supplementary figure 1**). Altogether, these results corroborate a central prediction of domain-general accounts of face recognition (Behrmann & Avidan, 2005; Garrido et al., 2018; Geskin & Behrmann, 2018; Grill-Spector et al., 2004; Kanwisher, 2000).

**Figure 2.**
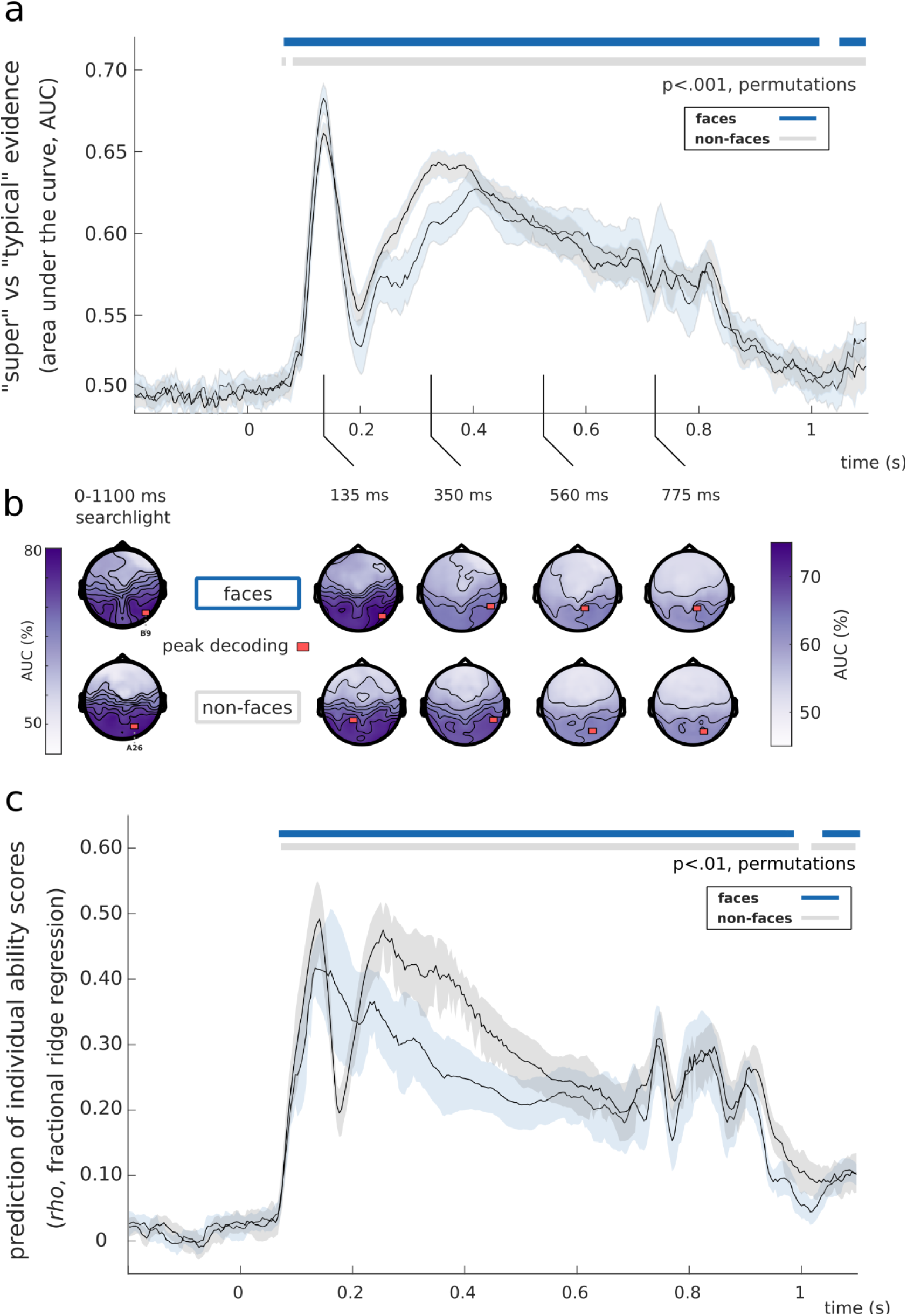
Decoding interindividual recognition ability variations from EEG activity. **a**) Trial-by-trial group-membership predictions (super-recogniser or typical recogniser) were computed from EEG patterns, for each 4-ms interval, while participants processed face (blue trace) or non-face stimuli (grey trace). Significant decoding performance occurred as early as 65 ms, peaked in the N170 window, and lasted for the remainder of the EEG epochs (*p*<.001). **b)** Topographies were obtained using searchlight decoding analyses, either concatenating all time points (left topographies) or for selected time-windows (right topographies). Concatenating all time points resulted in peak classification performance of 77.3% over right occipito-temporal electrodes for face and 77.5% over right occipito-temporal electrodes for non-face conditions. In the N170 window, we observed a peak classification performance of 74.8% over right-temporal electrodes for face, and 72.1% over left-temporal electrodes for non-face conditions. **c**) We decoded the CFMT+ scores of the typical recognisers using fractional ridge regression. This yielded similar results with significant decoding as early as 75 ms, peaking around the N170 time window (peak-*rho*_face_=.4149, peak-*rho*_non-face_=.4899), and lasted for the remainder of the EEG epochs (*p*<.01, 1K permutations, 10 repetitions).

### Predicting recognition ability from 1 second of brain activity

An ongoing debate in individual differences research is whether the observed effects emerge from qualitative or quantitative changes in the supporting brain mechanisms (Barton and Corrow 2016; Bobak et al. 2017; Rosenthal et al. 2017; Hendel et al. 2019; Vogel et al. 2005; Maguire et al. 2003; Zadelaar et al. 2019; Price and Friston 2002; Anderson et al. 2020). The decoding results presented up to this point might give the impression that face recognition ability is supported by qualitative differences in brain mechanisms. However, these results were obtained with dichotomous classification models applied, by design, to the brains of individuals from a bimodal distribution of ability scores (e.g. Maguire et al., 2003).

To better assess the nature of the relationship between neural representations and ability in the general population, we thus performed a new decoding analysis on the typical recognisers only, using a continuous regression model. Specifically, we used cross-validated fractional ridge regression (Rokem & Kay, 2020) to predict *individual* CFMT+ face recognition ability scores from single-trial EEG data. This showed essentially similar results to the previous dichotomic decoding results: performance was above statistical threshold (*p*<.01, FDR-corrected) from about 80 ms to at least 1 s, peaking around 135 ms following stimulus onset for both face and non-face stimuli (**Fig. 2b,** peak-rho_face_=.4149 at 133 ms, peak-rho_non-face_=.4899 at 141 ms). This accurate decoding of individual scores from EEG patterns is compatible with a quantitative account of variations in brain mechanisms across individuals differing in face recognition abilities. Altogether, these decoding results provide evidence for important, domain-general, quantitative and temporally extended variations in the brain activity supporting face recognition abilities. This extended decoding suggests effects of individual ability across multiple successive processing stages.

### Linking neural representations and computational models of vision

Decoding time courses, and evidence for domain generality, however, offer limited insights on the level of brain computations (Lamme & Roelfsema, 2000; McDermott et al., 2002). To better characterise the visual brain computations covarying with face recognition ability, we compared, using representational similarity analysis (Kriegeskorte, Mur, & Bandettini, 2008; Kriegeskorte, Mur, Ruff, et al., 2008; Kriegeskorte & Kievit, 2013; Charest et al., 2014), the brain representations of our participants to that of convolutional neural networks (CNNs) trained to categorise objects (Krizhevsky et al. 2012; Simonyan & Zisserman, 2014; Güçlü & van Gerven, 2015). These CNNs process visual features of gradually higher complexity and abstraction along their layers (Güçlü & van Gerven, 2015), from low-level (e.g., orientation, edges) to high-level features (e.g., objects and object parts).

The brain representations were characterised by computing representational dissimilarity matrices (RDMs) for each participant and for each 4-ms time interval. These brain RDMs were derived using the cross-validated decoding performance of a linear discriminant model, where brain activity was decoded for every pair of stimuli at a given time interval (Carlson et al., 2013; Cichy et al., 2014; see **Supplementary Fig. 2** for the group-average RDMs and time course of key categorical distinctions). The visual model representations were characterised by computing RDMs from the layers of the CNNs, using Pearson correlations of the unit activations across all pairs of stimuli. Compared to typical participants, we found that the brain RDMs of super-recognisers showed larger mutual information (Ince et al., 2017) with the layer RDMs of CNNs that represent mid-level features (e.g., combinations of edges, contour, shape, texture; Güçlü & van Gerven, 2015; Long et al., 2018) between 133 and 165 ms (**Fig. 3a**, *p*<.05, cluster-test; see also **Supplementary Fig. 3**).

**Figure 3.**
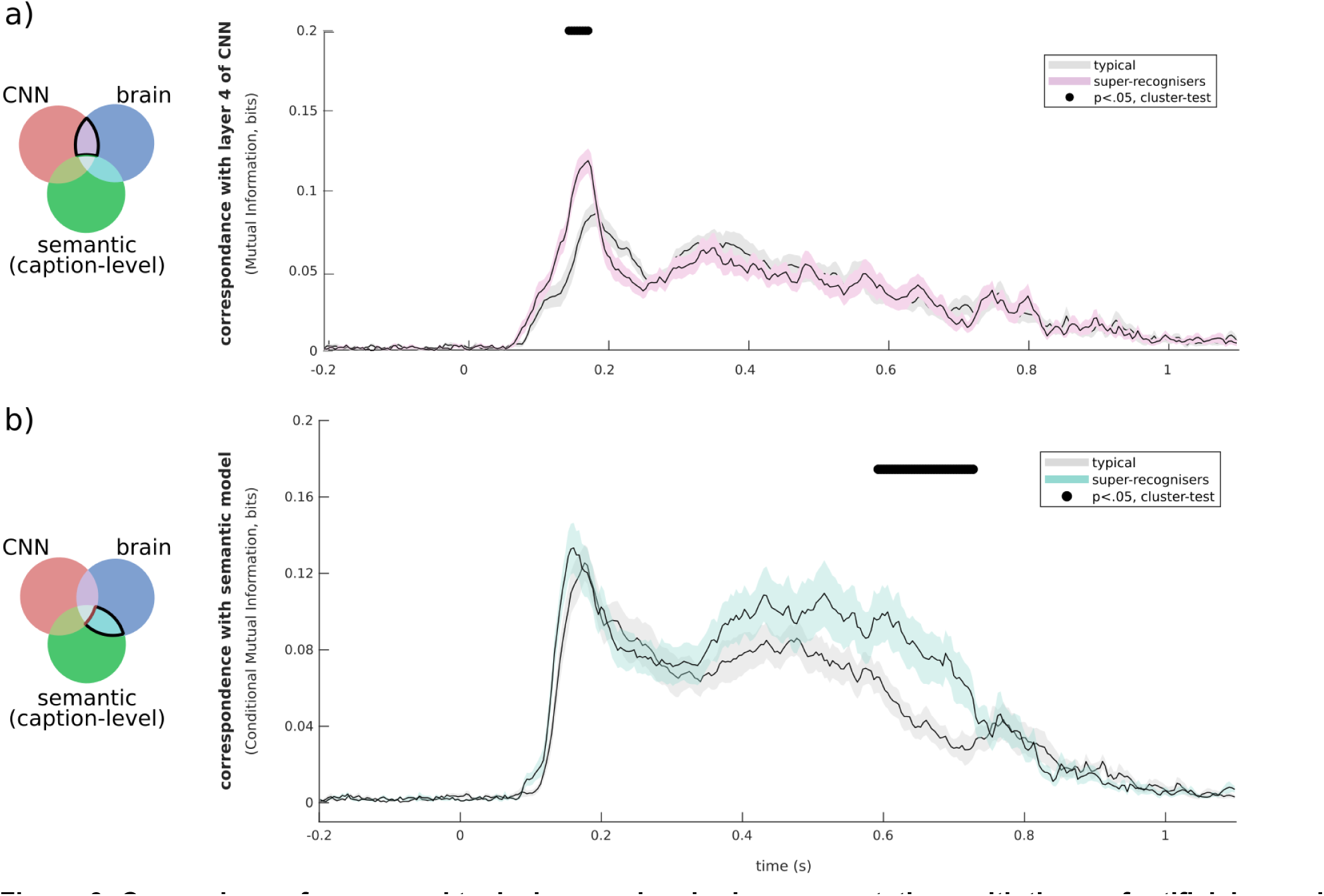
Comparison of super- and typical-recogniser brain representations with those of artificial neural networks of visual and semantic processing. **a**) Mutual information between brain RDMs and AlexNet RDMs (removing shared mutual information between brain and semantic model) is shown for typical- (grey curve) and super-recognisers (pink curve). We found greater similarity with mid-level visual computations (layer 4 shown, but similar results for layers 3 and 6 and for mid-layers of VGG16, another popular CNN model; see **Supplementary Fig. 3**) in the brains of super-recognisers (black line indicates significant contrasts, *p*<.05, cluster-corrected) between 133 ms and 165 ms. Similar results were observed when comparing brains and CNN models without removing the shared mutual information between brains and the semantic (caption-level) model (**Supplementary Fig. 3**). **b**) Mutual Information with the semantic model (excluding shared mutual information between brain and AlexNet) differed for typical- and super-recognisers in a later time window centred around 650 ms (cyan curve; super > typical, *p* < .05, cluster-corrected). Again, similar results were observed when comparing brains and the semantic model without removing the shared mutual information between the brain and AlexNet (**Supplementary Fig. 3**). The shaded areas of all curves represent the standard error.

### Linking neural representations with computational model of semantics

The finding that ability decoding was significant as late as 1 s after stimulus onset hints that brain computations beyond what is typically construed as pure visual processing also differ as a function of face recognition ability. To test this hypothesis, we asked five new participants to write captions describing the images presented during our experiment (e.g., “A city seen through a forest.”), and used a deep averaging network (Google Universal Sentence Encoder, GUSE; Cer et al., 2018) to transform these captions into embeddings (points in a caption space). GUSE has been trained to predict semantic textual similarity from human judgments, and its embeddings generalise to an array of other semantic judgement tasks (Cer et al., 2018). We then compared the RDMs computed from this semantic model to the brain RDMs of both typical- and super-recognisers. Importantly, both this comparison, and the one comparing brain and visual models, excluded the information shared between the semantic and visual models (but see **Supplementary Fig. 3** for similar results with unconstrained analyses). We found larger mutual information with these semantic computations in the brains of super-recognisers than in those of typical recognisers in a late window between 598 and 727 ms (**Fig. 3b**, *p*<.05, cluster-test).

### Comparing brain representations with those from human shape and semantic similarity judgements

Our findings so far suggest that mid-level visual and semantic brain processes both support individual differences in face recognition abilities. We further tested these conclusions using RDMs derived from a behavioural experiment on human participants. A group of 32 new participants were thus submitted to two multiple arrangement tasks engaging judgements of different levels of abstraction (Cichy et al. 2019; Mur et al. 2013; Hebart et al. 2018): in one of the tasks, they were asked to evaluate the shape similarities of the 49 object/face/scene images used in the main experiment; and, in the other task, they were instructed to judge the meaning similarities of the 49 sentence captions describing these images. More specifically, participants arranged the images/sentences on a computer screen inside a white circular arena according to the task instructions using simple drag and drop operations (see Fig. 4). We computed the mutual information between the mean RDMs extracted from each of these two tasks and the time-resolved brain RDMs of super- and typical recognisers, while excluding the information shared with the other task. Results indicated that shape representations were enhanced around mid-latencies in super-recognisers relative to typical recognisers (133-153 ms; *p*<.05, cluster-corrected; see Fig. 4a), while semantic meaning representations were enhanced in late latencies in super-recognisers compared to typical recognisers. (637-747 ms; *p*<.05, cluster-corrected; see Fig. 4b).

**Figure 4.**
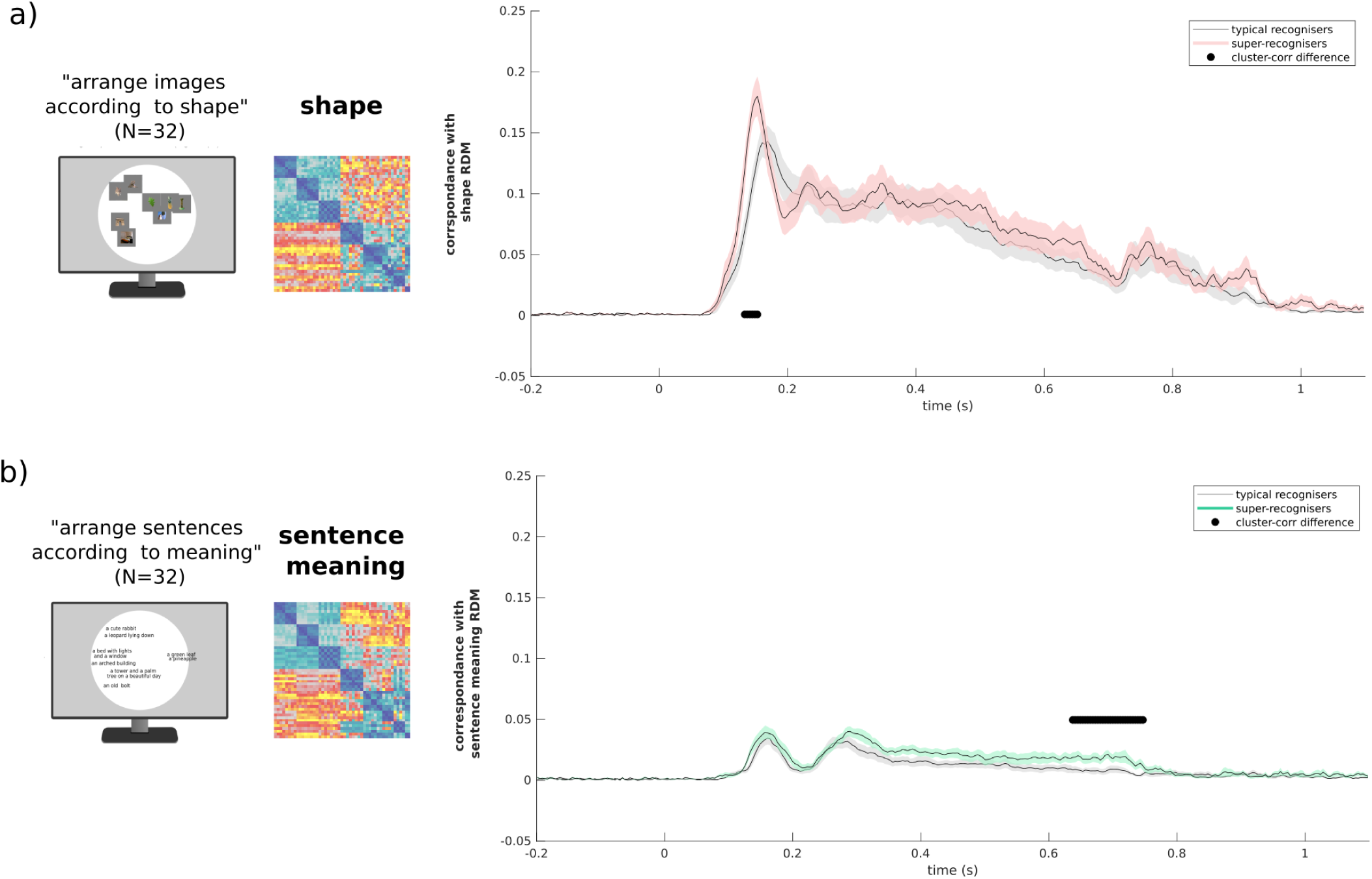
Comparison of brain representations with those from human shape and semantic similarity judgements. **a)** Mutual information between brain RDMs and the mean RDM built from shape similarity judgements (illustrated on the left of the plot), removing shared mutual information between brain and sentence caption meaning RDM, is shown for typical- (grey curve) and super-recognisers (red curve). We found greater similarity with shape information in the brains of super-recognisers between 133 ms and 153 ms (black line indicates significant contrasts, *p*<.05, cluster-corrected) — akin to the time interval during which we observed greater similarity between the brains of super-recognisers and mid-level layer CNN (fig. 3a). The shaded areas of all curves represent the standard error. **b)** Mutual information between brain RDMs and the mean RDM built from meaning similarity judgement of sentence captions (illustrated on the left of the plot), removing shared mutual information between brain and shape RDM, is shown for typical- (grey curve) and super-recognisers (green curve). We found greater similarity with sentence meaning in the brains of super-recognisers between 637 ms and 747 ms (black line indicates significant contrasts, *p*<.05, cluster-corrected), in agreement with our comparisons with the artificial semantic model (fig. 3b).

## Discussion

Using a data-driven approach combining neuroimaging, computational models, and behavioural tests, we characterised the computations modulated by variations in face recognition ability in the human brain. We recorded the high-density electroencephalographic (EEG) response to face and non-face stimuli in super-recognisers and typical recognisers. Using multivariate analysis, we reliably decoded group membership as well as recognition abilities of single individuals from a single second of brain activity. We then characterised the neural computations underlying these individual differences by comparing human brain activity with computations from artificial neural network models of vision and semantics using representational similarity analysis. Furthermore, we compared the representational geometries of these neural computations with those derived from additional human participants engaged in two tasks involving, respectively, shape similarity judgments on our stimuli and meaning similarity judgments on sentence captions describing these stimuli. These sets of comparisons revealed two main findings. First, we found higher similarity between early brain computations of super-recognisers and mid-level computations of vision models as well as shape similarity judgments. Second, this approach revealed higher similarity between late brain representations of super-recognisers and computations of an artificial semantic model as well as meaning similarity judgments. To our knowledge, this is the first demonstration of a link between face recognition ability and brain computations beyond high-level vision. Overall, these findings revealed specific computations supporting our individual ability to recognise faces, and suggest widespread variations in brain processes related to this crucial ability.

We achieved robust decoding of face recognition ability when examining EEG responses to face *and* non-face stimuli. This domain-general decoding result indicates that mechanisms underlying face recognition abilities give rise to enhanced neural representations that are not restricted to faces (Vinken et al. 2022). This is consistent with several neuropsychological (Barton et al., 2019; Bobak et al., 2016; Duchaine et al., 2007; Gabay et al., 2017; Geskin & Behrmann, 2018; Hendel et al., 2019) and brain imaging findings (Avidan et al., 2005; Jiahui et al., 2018; Kaltwasser et al., 2014; Rosenthal et al., 2017) showing face and non-face processing effects in individuals across the spectrum of face recognition ability (Behrmann & Plaut, 2013; Harel et al., 2013; but see Duchaine et al., 2006; Furl et al., 2011; Lohse et al., 2016; Wilmer et al., 2012). In addition, this decoding approach may provide quick and accurate alternatives to standardised behavioural tests assessing face recognition ability, for example in the context of security settings that benefit from strong face processing skills among their personnel (such as police agencies, border patrol, etc.). It could also be used in a closed-loop training procedure designed to improve face recognition ability (Faghel-Soubeyrand et al. 2019).

The decoding we observed for face and non-face stimuli peaked in the temporal window around the N170 component (Bentin et al., 1996). At that time, the computations in the brains of our participants differed most with respect to the mid-layer computations of artificial models of vision. These layers have been previously linked to processing in human infero-temporal cortex (hIT; Khaligh-Razavi and Kriegeskorte 2014; Güçlü and van Gerven 2015; Jiahui et al. ; Grossman et al. 2019) and functionally to mid-level feature representations such as combinations of edges and parts of objects (Güçlü & van Gerven, 2015; Long et al., 2018). We confirmed the visual nature of this representational code in another analysis associating brain to behaviour (human) shape representations. Such associations with the N170, however, does not mean that this component is exclusively involved in these mid-level processes. Rather, it suggests that other visual computations, including the high-level visual computations usually associated with the N170, do not differ substantially between super-recognisers and typical recognisers. The fact that these mid-level features are mostly shared between face and non-face stimuli could explain at least partly the high decoding performance observed for both classes of stimuli.

Finally, we found that face recognition ability is also associated with semantic computations that extend beyond basic-level visual categorisation in a late time-window around the P600 component (Eimer et al., 2012; Shen et al., 2016; van Herten et al., 2005). Recent studies using computational techniques have shown that word representations derived from models of natural language processing explain significant variance in the visual ventral stream (Popham et al. 2021; Dwivedi et al. 2021; Fernandino et al. 2022; Frisby et al. 2023). The current study goes beyond this recent work in two ways. First, our use of human sentence description and sentence encoders to characterise semantic (caption-level) computations provides a more abstract description of brain representations. Second, and most importantly, our work revealed a link between semantic brain computations and individual differences in face recognition ability. An association between semantic processes and face recognition ability had been posited in models of face recognition (Bruce & Young, 1986; Duchaine & Yovel, 2015) but, to our knowledge, it had never been shown empirically before.

With the development of novel and better artificial models simulating an increasing variety of cognitive processes, and with the technological advances allowing the processing of increasingly larger neuroimaging datasets, the approach described here provides a stepping stone for better understanding face recognition idiosyncrasies in the human brain.

## Methods

### Participants

A total of 33 participants were recruited for this study. The first group consisted of 16 individuals with exceptional ability in face recognition — super-recognisers. The second group was composed of 17 neurotypical controls. These sample sizes were chosen according to the effect sizes described in previous multivariate object recognition studies (Carlson et al., 2013; Cichy et al., 2014; Hebart et al., 2018). The data from one super-recogniser was excluded due to faulty EEG recordings. No participant had a history of psychiatric diagnostic or neurological disorder. All had normal or corrected to normal vision. This study was approved by the Ethics and Research Committee of the University of Birmingham, and informed consent was obtained from all participants.

Sixteen previously known super-recognisers were tested in the current study (30-44 years old, 10 female). Eight of these (SR1-SR8) were identified by Prof. Josh Davis from the University of Greenwich using an online test battery comprising a total of six face cognition tasks (Noyes et al., 2021) and tested at the University of Birmingham. The remaining eight (SR-9 to SR-16) were identified using three challenging face cognition tests (Ramon, 2021) and were tested at the University of Fribourg. The behavioural test scores for all participants are provided in **Supplementary Tables 1 and 2.** Across SR cohorts, the Cambridge Face Memory Test long-form (CFMT+; Russell et al., 2009) was used as the measure of face identity processing ability. A score greater than 90 (i.e., 2 SD above average) is typically considered the threshold for super-recognition (Bobak et al., 2016; Davis et al., 2016; Russell et al., 2009). Our 16 super-recognisers all scored above 92 (M=95.31; SD=2.68). A score of 92 corresponds to the 99^th^ percentile according to our estimation from a group of 332 participants from the general population recruited in three independent studies (Faghel-Soubeyrand et al., 2019; Fysh et al., 2020; Tardif et al., 2019).

An additional 17 typical recognisers (20-37 years old, 11 female) were recruited and tested on campus at the University of Fribourg (n=10) and the University of Birmingham (n=7). Their CFMT+ scores ranged from 50 to 85 (M=70.00; SD=9.08). Neither the average nor the distribution of this sample differed significantly from those of the 332 participants from the general population mentioned above (see **Fig. 1a**; t(346)=1.3065, *p*=0.1922; two-sample Kolmogorov-Smirnov test; D(346)=0.2545, *p*=0.2372).

### Tasks

#### CFMT+

All participants were administered the CFMT long-form, or CFMT+ (Russell et al., 2009). In the CFMT+, participants are required to memorise a series of face identities, and to subsequently identify the newly learned faces among three faces. It includes a total of 102 trials of increasing difficulty. The duration of this test is about 15 minutes. EEG was not recorded while participants completed this test.

#### One-back task

##### Stimuli

The stimuli used in this study consisted of 49 images of faces, animals (e.g., giraffe, monkey, puppy), plants, objects (e.g., car, computer monitor, flower, banana), and scenes (e.g., city landscape, kitchen, bedroom). The 24 faces (13 identities, 8 males, and 8 neutral, 8 happy, 8 fearful expressions) were sampled from the Radboud Face dataset (Langner et al., 2010). The main facial features were aligned across faces using Procrustes transformations. Each face image was revealed through an ellipsoid mask that excluded non-facial cues. The non-face images were sampled from the stimulus set of Kiani et al. (Kiani et al., 2007). All stimuli were converted to 250 x 250 pixels (8x8 deg of visual angle) greyscale images. The mean luminance and the luminance standard deviation of these stimuli were equalised using the SHINE toolbox (Willenbockel et al., 2010).

##### Procedure

We measured high-density electroencephalographic (EEG; sampling rate = 1024 Hz; 128-channel BioSemi ActiveTwo headset) activity while participants performed ∼3200 trials of a one-back task in two recording sessions separated by at least one day and by a maximum of two weeks (**Fig. 1b**). Participants were asked to press a computer keyboard key on trials where the current image was identical to the previous one. Repetitions occurred with a 0.1 probability. They were asked to respond as quickly and accurately as possible. Feedback about accuracy was given on each trial. A trial unravelled as follows: a white fixation point was presented on a grey background for 500 ms (with a jitter of ± 50 ms); followed by a stimulus presented on a grey background for 600 ms; and, finally, by a white fixation point on a grey background for 500 ms. Participants had a maximum of 1100 ms following stimulus onset to respond. This interval, as well as the 200 ms preceding stimulus onset, constituted the epoch selected for our EEG analyses. In total, our participants completed 105,600 one-back trials which constituted ∼32 hours of EEG epochs.

##### Shape and sentence meaning multiple arrangements tasks

Thirty four new neurotypical participants took part in two multiple arrangements tasks (e.g. Kriegeskorte and Mur, 2012; Charest et al., 2014) in counterbalanced orders. In one of the tasks, they were asked to evaluate the shape similarities of the 49 stimuli used in the main experiment while, in the other task, they were instructed to judge the meaning similarities of sentence captions describing these stimuli (see Semantic Caption-level Deep Averaging Neural Network RDM for more information about these sentence captions).

More specifically, participants were asked to arrange stimuli or sentence captions on a computer screen inside a white circular arena by using computer mouse drag and drop operations. During the shape (vs. meaning) multiple arrangement task, they were instructed to place the displayed visual stimuli (vs. sentence captions) in such a way that their pairwise distances match their shape (vs. meaning) similarities as much as possible (Fig. 4). On the first trial of each task, participants arranged all 49 items. On subsequent trials, a subset of these items was selected based on an adaptive procedure aimed at minimising uncertainty for all possible pairs of items (e.g. items that initially were placed very close to each other) and at better approximating the high-dimensional perceptual representational space (Kriegeskorte and Mur, 2012). This procedure was repeated until the task timed out (20 min).

We computed one RDM per task per participant. Two participants were excluded from the final sample because their RDMs differed from the mean RDMs by more than two standard deviations. Finally, we averaged the remaining individual RDMs within each task.

### Analyses

All reported analyses were performed independently for each EEG recording session and then averaged. Analyses were completed using custom code written in MATLAB (MathWorks) and Python.

### EEG preprocessing

EEG data was preprocessed using FieldTrip (Oostenveld et al., 2011): continuous raw data was first re-referenced relative to Cz, filtered with a band-pass filter [0.01-80 Hz], segmented into trial epochs from −200 ms to 1100 ms relative to stimulus onset, and down-sampled at 256 Hz.

### Decoding analyses

#### Whole-brain analyses

To predict group-membership from EEG brain activity, we trained Fisher linear discriminant classifiers (5-fold cross-validation, 5 repetitions; Treder 2020; Grootswagers et al. 2017), using all 128 channels single-trial EEG data as features. Separate analyses were done on either all time points after stimulus onset, or successive 4-ms EEG time intervals. The classifiers were trained on trials of EEG activity of participants viewing faces and non-faces independently. The Area Under the Curve (AUC) was used to assess sensitivity. Additional control decoding analyses investigating effects of one-back trials on the predictions are shown in **Supplementary Fig. 1.**

#### Searchlight analysis

We conducted a searchlight analysis decoding EEG signals from all subsets of five neighbouring channels to characterise the scalp topographies of group-membership AUC. This searchlight analysis was done either using the entire EEG time series of a trial (0-1100 ms; **Fig. 2b,** leftmost topographies), or using 60 ms temporal windows (centred on 135 ms, 350 ms, 560 ms, and 775 ms; **Fig. 2b** rightmost topographies). We ran additional control searchlight decoding procedures investigating the effect of one-back trials (**Supplementary Fig. 1**).

#### Regression analysis

We used fractional ridge regression models (Rokem & Kay, 2020) to predict individual face recognition ability scores (CFMT+) among the typical recognisers from EEG patterns across time. We trained our model on subsets of 60% of the EEG patterns. We chose the alpha hyperparameter with the best coefficient of determination among 20 alpha hyperparameters ranging linearly from 0.001 to 0.99 applied on a 30% validation set. The decoding performance was assessed using the Spearman correlation between the CFMT+ scores and predictions from the overall best model (applied on the remaining 10% of EEG patterns). This process was repeated 10 times and the Spearman correlations were averaged. Significance was assessed using a permutation test (see *Group comparisons and inferential statistics* section).

### Representational Similarity Analysis of brain and computational models

We compared our participants’ brain representations to those from visual and semantic (caption-level) artificial neural networks using Representational Similarity Analysis (RSA; Charest et al., 2014; Kriegeskorte, Mur, & Bandettini, 2008; Kriegeskorte, Mur, Ruff, et al., 2008; Kriegeskorte & Kievit, 8/2013).

#### Brain Representational Dissimilarity Matrices

For every participant, we trained a Fisher linear discriminant to distinguish pairs of stimuli from every 4-ms intervals of EEG response (on all 128 channels) to these stimuli from −200 to 1100 ms after stimulus onset (Cichy & Oliva, 2020; Graumann et al., 2022). Cross-validated AUC served as pairwise classification dissimilarity metric. By repeating this process for all possible pairs (1176 for our 49 stimuli), we obtained a representational dissimilarity matrix (RDM). RDMs are shown for selected time points in **Supplementary Fig. 2**.

#### Visual Convolutional Neural Networks RDMs

We used a pre-trained AlexNet (Krizhevsky et al., 2012) as one model of the visual computations along the ventral stream (Güçlü & van Gerven, 2015). Our 49 stimuli were input to AlexNet. Layer-wise RDMs were constructed comparing the unit activation patterns for each pair of images using Pearson correlations. Similarly, we computed layer-wise RDMs from another well-known CNN, VGG-16 (see **Supplementary Fig. 3b**). Following previous studies using this model (Liu et al. 2021; Xie et al. 2020), we averaged the convolutional layer RDMs situated between each max pooling layers and the layers’ input into five aggregated convolutional RDMs (e.g. conv1-1 & conv1-2 into RDM-conv1); this facilitated the comparison of our results with the five convolutional layers of AlexNet.

#### Semantic Caption-level Deep Averaging Neural Network RDM

We asked five new participants to provide a sentence caption describing each stimulus (e.g., “a city seen from the other side of the forest”, see **Fig. 1d**) using the Meadows online platform (www.meadows-research.com). The sentence captions were input in Google’s universal sentence encoder (GUSE; Cer et al., 2018) resulting in 512 dimensional sentence embeddings for each stimulus. We then computed the dissimilarities (cosine distances) between the sentence embeddings across all pairs of captions, resulting in a semantic caption-level RDM for each participant. The average RDM was used for further analyses.

### Comparing brain representations with computational models

We compared our participants’ brain RDMs to those from the vision (Fig. 3a) and semantic (Fig. 3b) models described in the previous section using Conditional Mutual Information (CMI, Ince et al., 2017), which measures the statistical dependence between two variables (e.g. mutual information *I*(x;y)), removing the effect from a third variable (i.e. I(x;y|z)). Additional control comparisons using unconstrained Mutual Information between brain RDMs and both models are shown in **Supplementary Fig. 3a.**

### Group comparison and inferential statistics

#### Comparison of Conditional Mutual Information time courses

Time courses of CMI were compared between the super-recognisers and typical recognisers using independent samples t-tests and a Monte Carlo procedure at a *p-value* of .05, as implemented in the Fieldtrip Toolbox (Oostenveld et al., 2011). Family-wise errors were controlled for using cluster-based corrections, with maximum cluster size as cluster-level statistic and an arbitrary *t* threshold for cluster statistic of [-1.90, 1.90] for the comparison of brain and semantic (excluding CNN) and [-2.75 2.75] for the comparison of brain and CNN (excluding semantic) time courses. The standard error is shown for all curves as colour-shaded areas (**Fig. 3**). Analyses with MI (brain; CNN) and MI (brain; semantic) were completed in an identical manner (**Supplementary Fig. 3**).

#### Time course of group-membership decoding

Significance was assessed using non-parametric permutation tests. We simulated the null hypothesis by training the linear classifier to identify shuffled group-membership labels from the experimental EEG patterns. This process was repeated 1000 times for each time point and each one of the two sessions. We then compared the real, experimental decoding value at each time point to its corresponding null distribution, and rejected the null hypothesis if the decoding value was greater than the prescribed critical value at a *p* <.001 level.

#### Time course of individual ability decoding using ridge regression

Significance was again assessed using non-parametric permutation testing. The ridge regression analysis predicted cross-validated CFMT+ scores from single trial EEG patterns, and goodness of fit is reported using Spearman’s correlation between the predicted and observed CFMT+ scores. Under the null hypothesis that all participants elicited comparable EEG response patterns, irrespective of their CFMT+ score, the face recognition ability scores are exchangeable. We simulated this null hypothesis by repeating the ridge regression model training using randomly shuffled CFMT+ scores. The predicted CFMT+ scores were then correlated to the empirical, observed CFMT+ scores using Spearman’s correlation, and this was repeated 1000 times for each time point. We finally compared the real, experimental correlation value with its corresponding null distribution at each time point, and rejected the null hypothesis if the correlation value was greater than the prescribed critical value at a *p* <.01 level.

## Data availability

Data associated with this article will be available online upon publication of the manuscript.

## Code availability

The MATLAB and Python codes used in this study will be available online upon publication of the manuscript.

## Acknowledgements

We thank Prof. Josh P. Davis for sharing behavioural scores of super-recognisers and establishing first contact to the UK-based Super-Recognizers reported here. Funding for this project was supported by an ERC Starting Grant [ERC-StG-759432] to I.C, an ERSC-IAA grant to J.W., I.C. and S.F.S., by a Swiss National Science Foundation PRIMA (Promoting Women in Academia) grant [PR00P1_179872] to MR, and by NSERC and IVADO graduate scholarships to S.F.S. We also thank Mick Neville, from Super-Recognisers Ltd., who helped us to get in contact with some of our super-recognizer participants. We thank Rose Jutras, who helped with data acquisition.

## Author contributions

**(CRediT standardised author statement)**

**S.F-S.**: Conceptualisation, methodology, software, formal analysis, investigation, data curation, writing - original draft, visualisation, supervision, project administration, funding acquisition. **M.R.:** Investigation, resources, project administration, writing - review and editing. **E.B.:** investigation, project administration. **M.Z.**: investigation. **J.W.:** funding acquisition, writing - review and editing. **A-R.R.**: Investigation. **R.C.:** Resources. **F.G.:** Methodology, writing - original draft, supervision, funding acquisition. **I.C.**: Supervision, methodology, software, resources, formal analysis, writing - original draft, project administration, funding acquisition.

## Supplementary material

### Behavioural results

All participants’ face recognition ability was assessed using the Cambridge Face Memory Test long-form (CFMT+, <u>(Russell et al. 2009)</u>). Scores on the CFMT+ ranged from 50 to 85 in the typical recognisers group (M_TRs_=70.00; SD=9.09), and from 92 to 100 in the experimental super-recogniser group (M_SRs_=95.38, SD=2.68; difference between groups : *t*(31)=10.6958, p<.00001 see Fig. 1a). The main experimental task was a one-back task (Fig. 1b). Accuracy was significantly greater for the super-recognisers (M_SRs_=.93, SD=.054) than for the typical recognisers (M_TRs_=.83, SD=.094; *t*(31)=3.7911, *p*=0.00065). This was also true when analysing separately face (M_SRs_= .9260, SD=.0471; M_TRs_=.8144, SD=.1066; *t*(31)=3.8440, *p*=0.00056) and non-face trials (M_SRs_=.9456, SD=.0651; M_TRs_=.8532, SD=.0929; *t*(31)=3.2855, *p*=.0025).

Furthermore, accuracy in the one-back task was positively correlated with scores on the CFMT+ (r=.68, p<.001; RT was marginally associated with CFMT+, *r*=.37, p=.04). We observed no significant differences in response times between the two groups (p>.3).

### Scores obtained by the super-recogniser tested in the UK on a battery of face recognition tests

**Supplementary table 1:** The scores above show performance on a battery of standardised face recognition tests (Noyes et al. 2021) for the participants identified as super-recognisers in the UK. Also shown are the scores for our one-back task, for face and non-face trials. The last row of the table shows the maximum obtainable score for each test.

| subject-ID | CFMT+ | GFMT | Face Array | LASIE Match | Black and White | Super-rec | Longterm | one-back faces | one-back non-faces |
| --- | --- | --- | --- | --- | --- | --- | --- | --- | --- |
| SR-1 | 93 | 40 | 38 | 83 | 34 | 13 | 9 | .9328 | 0.977 |
| SR-2 | 95 | 40 | 37 | 88 | 33 | 12 | 9 | .8500 | 0.7903 |
| SR-3 | 97 | 40 | 32 | 84 | 33 | 12 | 8 | .9325 | 0.9718 |
| SR-4 | 93 | 40 | 38 | 92 | 34 | 13 | 8 | .9726 | 0.9835 |
| SR-5 | 98 | 40 | 31 | 91 | 33 | 14 | 9 | .9326 | 0.9787 |
| SR-6 | 100 | 40 | 39 | 82 | 40 | 14 | 10 | .9823 | 0.9817 |
| SR-7 | 98 | 40 | 40 | 92 | 33 | 13 | 8 | .9319 | 0.9647 |
| SR-8 | 96 | 38 | 40 | 90 | 39 | 12 | 10 | .9362 | 0.9787 |
| Max score | 102 | 40 | 40 | 100 | 40 | 14 | 10 | 1 | 1 |

### Scores obtained by the super-recogniser tested in Switzerland on a battery of face recognition tests

**Supplementary table 2:** The scores above show performance on a battery of standardised face recognition tests (Ramon, 2021) for the participants identified as super-recognisers in Switzerland. Also shown are the scores for our one-back task, for face and non-face trials. The last row of the table shows the best obtainable score for each test.

| subject-ID | CFMT+ | FICST score | YBT long raw score | one-back faces | one-back non-faces |
| --- | --- | --- | --- | --- | --- |
| SR-9 | 92 | 0 | 29 | .8132 | 0.8384 |
| SR-10 | 99 | 0 | 17 | .9409 | 0.9619 |
| SR-11 | 92 | 0 | 20 | .9635 | 0.9808 |
| SR-12 | 93 | 1 | 20 | .8581 | 0.8212 |
| SR-13 | 92 | 7 | 17 | .9530 | 0.985 |
| SR-14 | 96 | 0 | 18 | .9604 | 0.9886 |
| SR-15 | 94 | 3 | 16 | .9112 | 0.9624 |
| SR-16 | 97 | 1 | 22 | .9792 | 0.9643 |
| Best score | 102 | 0 | 35 | 1 | 1 |

### Noise ceiling for group-membership decoding using CFMT+ reliability scores

To better interpret the magnitude of our group-membership decoding accuracies, we determined the maximum attainable accuracy when categorising a super-recogniser as such using the CFMT+ (i.e., the empirically imposed noise ceiling for our decoding group-membership analysis). We simulated 1000 distributions of CFMT+ scores, each with N=32, and each having a Pearson correlation of .71±.01 with the distribution of CFMT+ scores observed in this study. The correlation coefficient of .71 was chosen according to the test-retest reliability the CFMT+ measured in previous studies (Arrington et al., 2022; Murray & Bate, 2020). We then predicted super-recogniser participants from these distributions: we compared the simulated SR label (i.e. simulated CFMT+ scores higher than the prescribed cut-off of 92) to the used SR labels. We then averaged the accuracies across the simulated participants, and computed the maximum accuracy from these 1000 distributions. This process was repeated 10000 times, creating a distribution of 10000 simulated maximums with a mean of M=0.9310 (SD=0.0228) that we interpreted as the noise ceiling.

### Univariate analyses of EEG associated with individual ability

#### N170 amplitude and latency

We also performed more traditional event-related potential (ERP) analyses for both groups. We extracted, for face and non-face trials, peak negative ERP amplitudes and latencies for every participant in a window corresponding to the N170 component (within 110-200 ms at electrodes [B6, B7, B8, A28] on the right hemisphere and [A9, A10, A11, A15] on the left hemisphere). We tested the conditions, groups, and their interaction effects using an ANOVA on N170 peak latency and amplitude separately. No interaction effects were observed for peak latencies (F_interaction_(60,1)=1, p=.32) and amplitudes (F_interaction_(60,1)=0.32, p>.50). The peak N170 was earlier (F_conditions_(60,1)=5.86, p=.0185) and presented greater amplitudes (F_conditions_(60,1)=33.78, p<.0001) for faces compared to non-face objects. Moreover, the peak N170 was earlier (F_group_(60,1)=19.23, p<.0001) and presented larger amplitudes (F_group_(60,1)=13.75, p=.0005) in super-recognisers than typical recognisers.

#### Lateralisation

Compared to typical recognisers, super-recognisers showed greater N170 peak amplitudes in the right-hemisphere electrodes for faces (computed as the difference between the right [B6, B7, B8, A28] and left [A9, A10, A11, A15]; t(30)=-2.8542, p=.01). Moreover, the CFMT+ scores of typical recognisers correlated with right-hemisphere lateralisation of the N170 for faces (r(16)=-.53, p=.0298). These effects were not significant either for non-face stimuli, or for N170 peak latency.

**Supplementary figure 1.**
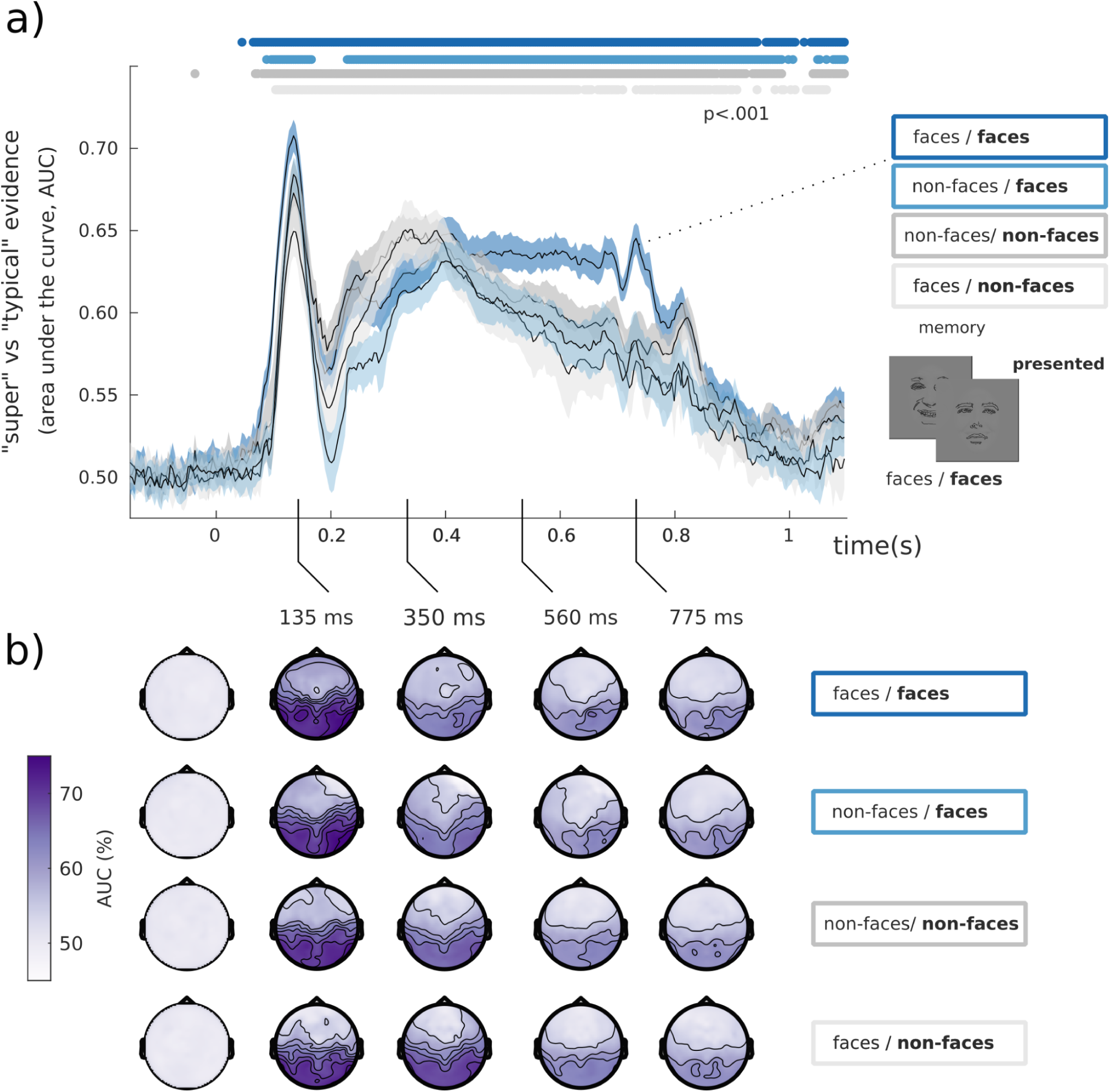
a) We computed the time course of decoding accuracy for group membership from all different face and non-face combinations of one-back and current trials (e.g. consecutive face - face trials). We observed a similar time course for all combinations. However the consecutive face-face trials showed larger decoding accuracies around 400 to 750 ms. b) The topographies show results from a searchlight decoding analysis with classification performance attaining 75% accuracy around 135 ms over occipito-temporal electrodes for face presented conditions (74.6% for face-face, 75.1% for nonface-face) and 72% for non-face presented conditions (71.7% for nonface-nonface, 71.5% for face-non-face). Note that drawings of faces are depicted here as an anonymised substitute to the experimental face stimuli presented to our participants.

**Supplementary figure 2.**
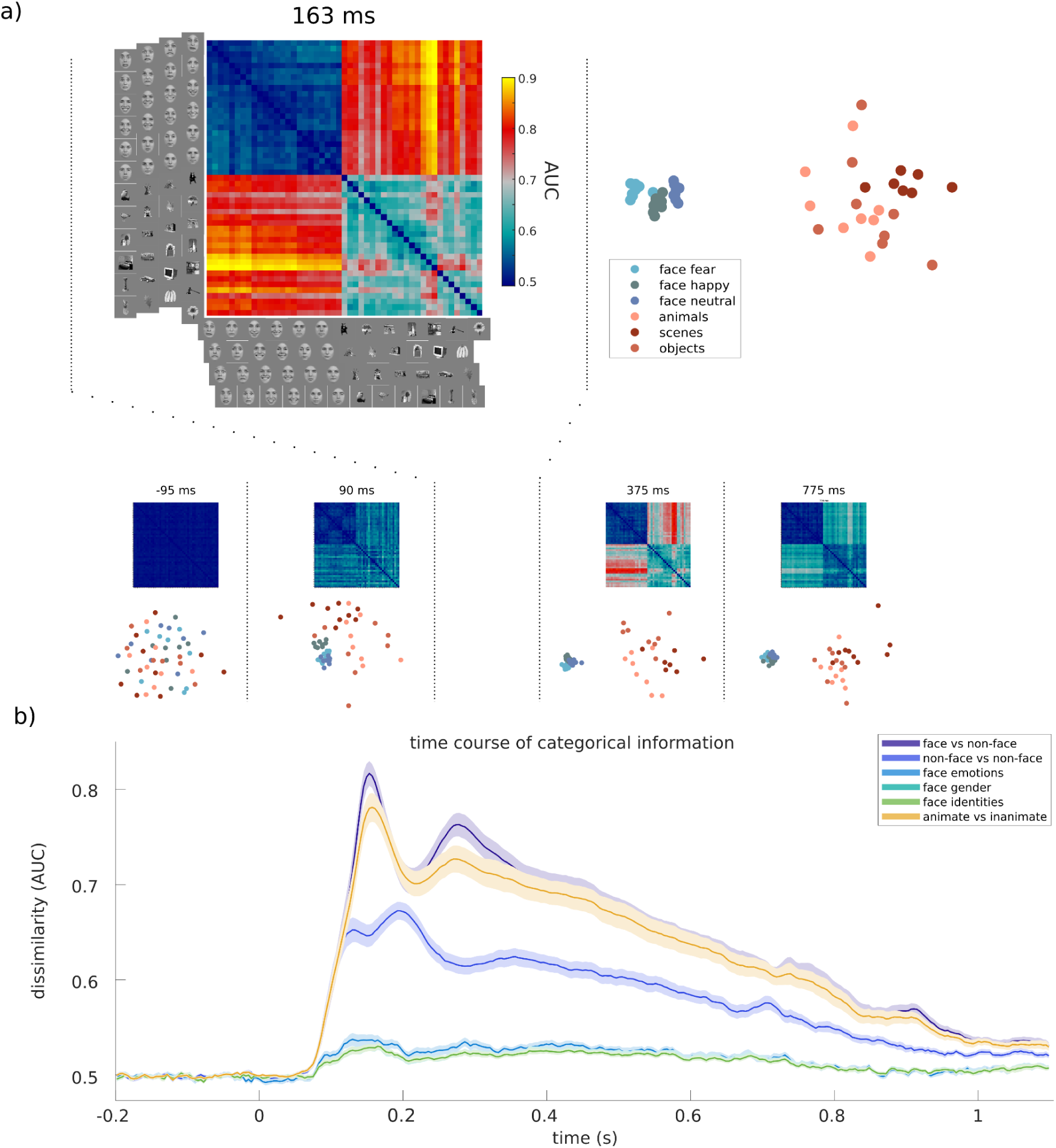
EEG representational geometry dynamics. a) Representational Similarity Analysis (RSA) was applied to time-resolved EEG patterns, using decoding AUC as dissimilarity measure between pairs of images (5 fold cross-validation, 5 repetitions) to create Representational Dissimilarity Matrices (RDMs). Multidimensional scaling was employed to visualise these high-dimensional brain representations on a 2D plane, which showed clear distinctions between various categories (e.g. face clusters, scenes clusters, animal clusters, etc.). b) We revealed categorical information unfolding in time by averaging dissimilarities between stimulus categories (e.g. faces vs non-face objects) and averaging across participants. Brain representations for the distinction of face vs. non-face objects (a hallmark of the N170, (Rossion & Jacques, 2012)) dominated all other categorical distinctions (Carlson et al., 2013; Kaneshiro et al., 2015), and peaked at 153 ms.

**Supplementary figure 3.**
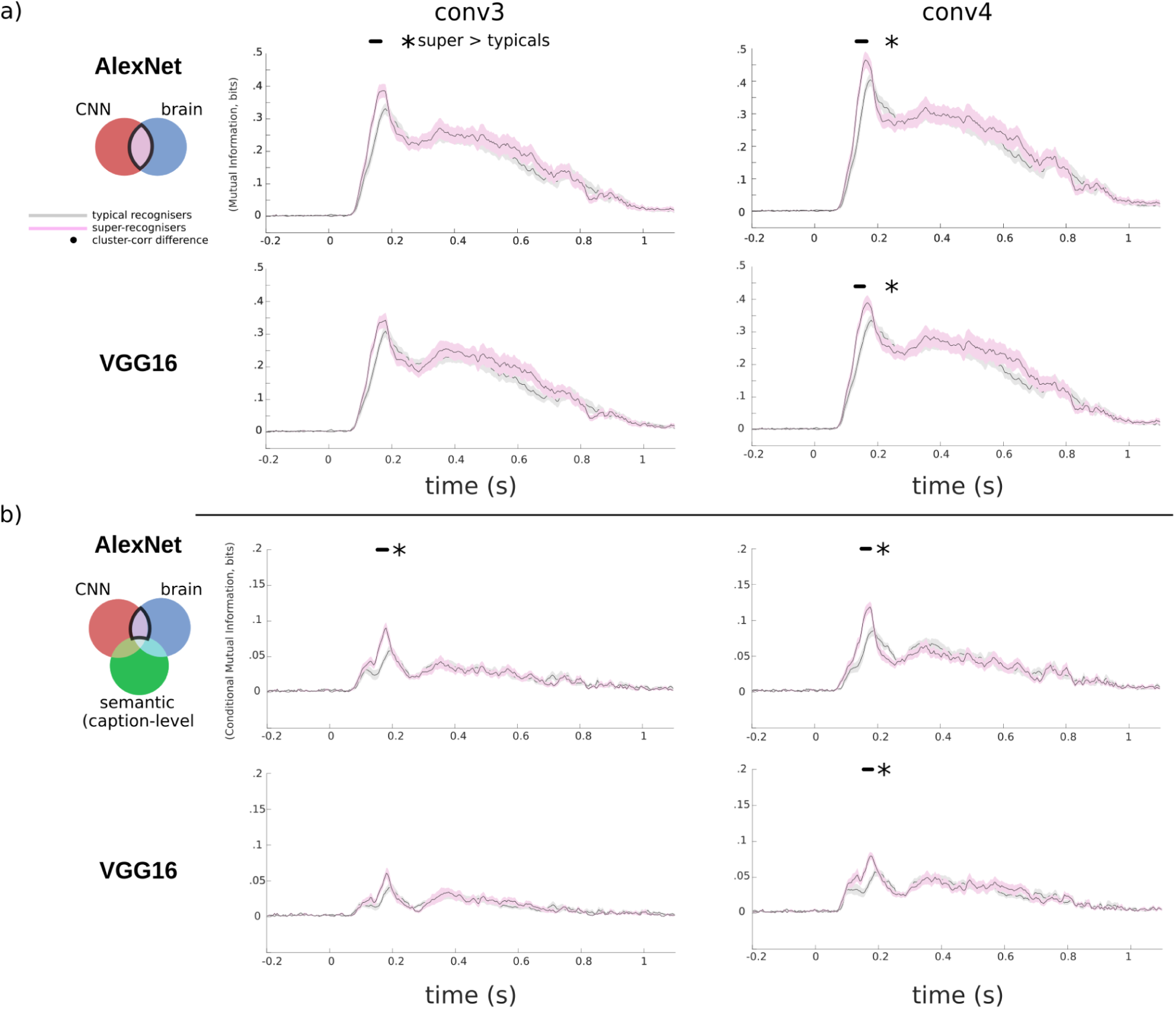
Comparison of super- and typical-recogniser brain representations with those of artificial neural networks of vision. **a**) Mutual information results comparing brain RDMs and AlexNet RDMs (first row) and brain RDMs and VGG16 RDMs (second row) are shown for typical- (grey curve) and super-recognisers (pink curve). We found greater similarity with mid-level visual computations as indexed from both CNN models (layers 3, 4 shown for AlexNet and VGG 16) in the brains of super-recognisers (black line indicates significant contrasts, *p*<.05, cluster-corrected) between 130 ms to 160 ms. **b**) We also computed the MI between CNN RDMs and the brain RDMs, but constrained on the Mutual Information from the caption-level semantic RDM. Again, we found greater similarity with mid-level visual computations as indexed from both CNN models (layers 3, 4 shown for AlexNet and VGG 16) in the brains of super-recognisers *(p*<.05, cluster-corrected) between 133 ms to 165 ms. The observed magnitudes were reduced as expected given the additional constraints. The shaded areas of all curves represent the standard error.

